# Reconstructing Phylogenetic Relationships Based on Repeat Sequence Similarities

**DOI:** 10.1101/624064

**Authors:** Daniel Vitales, Sònia Garcia, Steven Dodsworth

## Abstract

A recent phylogenetic method based on genome-wide abundance of different repeat types proved to be useful in reconstructing the evolutionary history of several plant and animal groups. Here, we demonstrate that an alternative information source from the repeatome can also be employed to infer phylogenetic relationships among taxa. Specifically, this novel approach makes use of the repeat sequence similarity matrices obtained from the comparative clustering analyses of RepeatExplorer 2, which are subsequently transformed to between-taxa distance matrices. These pairwise matrices are used to construct neighbour-joining trees for each of the top most-abundant clusters and they are finally summarized in a consensus network. This methodology was tested on three groups of angiosperms and one group of insects, resulting in congruent evolutionary hypotheses compared to more standard systematic analyses based on commonly used DNA markers. We propose that the combined application of these phylogenetic approaches based on repeat abundances and repeat sequence similarities could be helpful to understand mechanisms governing genome and repeatome evolution.

## Introduction

Repetitive elements make up the major proportion of all the nuclear DNA in most eukaryotic genomes (Biscotti et al. 2015), contributing up to 70-80% of the genome in the case of angiosperms (Kelly et al. 2012). The repetitive fraction of the genome is also very diverse, showing large variability across taxa in terms of structure (e.g. Neumann et al. 2019), abundance (e.g. Staton & Burke 2015) and chromosomal position (e.g. Richard et al. 2008) of different classes of tandem repeats and interspersed mobile elements (i.e. DNA transposons and retrotransposons). This diversity has largely been studied with the use of classical techniques like fluorescence in-situ hybridization, PCR, cloning, Sanger sequencing or blot methods, positively contributing to our understanding of organismal evolutionary history (Weiss-Schneeweiss et al. 2015). However, mainly due to sequencing and analytical difficulties inherent to the repetitive nature of this component of the genome, systematic researchers have largely avoided the use of the repeatome as a source of phylogenetic markers.

The arrival of high-throughput sequencing (HTS) deeply changed the possibilities to investigate repetitive elements. These new technologies are a fast and inexpensive way of gathering sequence data from repeats across the entire genome (Weiss-Schneeweiss et al. 2015). In parallel with technical advances in DNA sequencing, the development of new bioinformatics tools has greatly facilitated the study of repetitive elements. Previous studies have shown that genome skimming followed by graph-based clustering of short sequence reads, is an easy and cost-effective combination to characterize the repeatome of species *de novo* (Macas et al. 2007; Novak et al. 2010). Taking advantage of this progress, Dodsworth et al. (2014) developed a new phylogenetic methodology based on the abundance of different repetitive elements. Briefly, their approach utilizes comparative clustering of HTS short reads to estimate abundances of repetitive DNAs on different species. Then, assuming that repeat amounts evolve primarily by random genetic drift (Jurka et al. 2011; 2012), their relative abundances are analysed as continuous characters for phylogenetic inference. The method proved to be useful in reconstructing the evolutionary history of several plant and animal groups (e.g. Dodsworth et al. 2014; Martín-Peciña et al. 2018; Bolsheva et al. 2019), yielding as well additional data for comparative studies of genome evolution. More recently, other authors have also proposed more or less analogous phylogenetic approaches based on the genomic abundance of repetitive elements (Piednöel et al. 2015; Harkess et al. 2016; Doronina et al. 2017), which emphasize the interest of evolutionary researchers on this topic.

In the present study, we use a different information source from the repeatome to reconstruct phylogenetic relationships among taxa. Specifically, this novel approach is based on the sequence similarities between homologous repetitive elements present in closely-related taxa (Fig. 1). As in the previous method based on repeat abundances (i.e. Dodsworth et al. 2014), the identification of homologous repeat classes first obtained from comparative graph-based clustering (Novak et al. 2010) is also a key step of the new approach (Fig. 1b). By running a comparative clustering analysis, combined sequence reads from different taxa are subjected to pairwise (BLAST) comparison. The latest implementation of the graph-based clustering pipeline [i.e. RepeatExplorer2 (Novak et al. 2013)] generates a similarity matrix – counting the number of reads of one taxon showing significant similarity hits (i.e. 90% similarity over at least 55% of their length) with reads from other taxa – for each of the most abundant estimated repeat families (clusters). In addition, by taking into account the number of reads of each species that constitutes the cluster, a derived matrix displays the observed/expected inter-specific read similarity values (Fig. 1c). This second matrix reports the pairwise sequence similarity for each repeat class, irrespectively of the number of reads for each species included in the comparative clustering analysis. Given that lineage-specific differences among repeats are mainly determined by diversification processes (Jurka et al. 2011; Dodsworth et al. 2014), the similarity matrices derived from each of the repeat clusters should contain a demonstrable phylogenetic signal.

**Figure 1.**
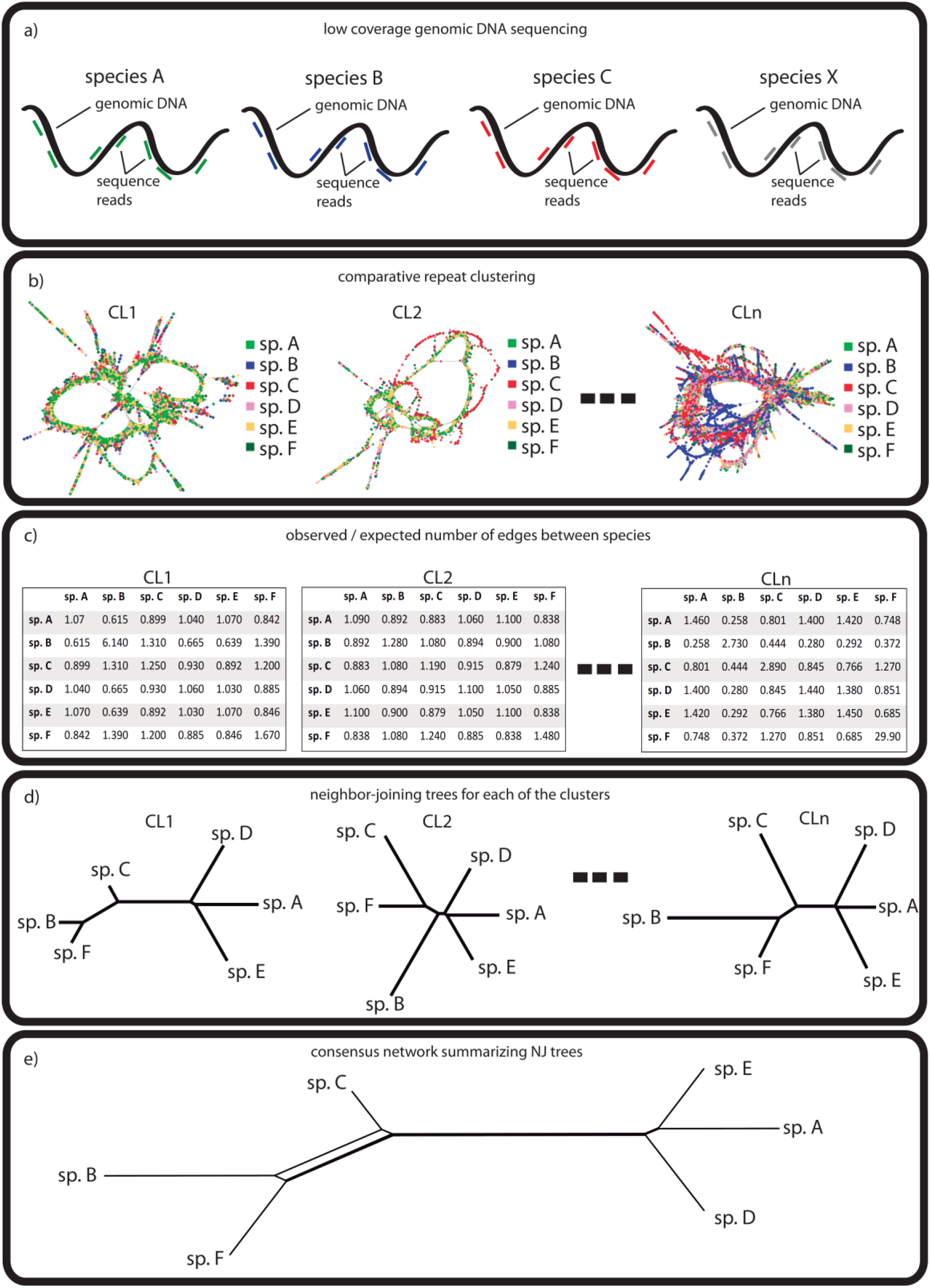
Schematic illustration of the workflow used for building trees from repeat sequence similarities. a) low-coverage genomic DNA sequencing using next-generation sequencing methods (NGS; e.g. Illumina). b) comparative clustering of NGS reads using RepeatExplorer pipeline. c) similarity matrices with the observed/expected number of edges between species. d) construction of NJ trees based on pairwise distance matrices from each cluster. e) consensus network summarizing the information of NJ trees.

To investigate this method, we chose four groups of organisms that had already been studied using the repeat abundance approach to infer phylogenetic hypotheses (Dodsworth et al. 2014). These study cases represent diverse evolutionary groups (three orders of angiosperms and one insect group) showing a variety of systematic problems and taxonomic levels. For all of them, we inferred the phylogenetic relationships among the species using their differential repeat sequence similarities. Subsequently, we tested the level of congruence of the new approach with previous evolutionary reconstructions based on repeat abundances and other more standard phylogenetic markers. Finally, we evaluated the performance of the method under different analysis configurations.

## Materials and methods

### Plant material and genomic data of study cases

#### Nicotiana

*Nicotiana* sect. *Repandae* is a group of four allotetraploid species originating from a single allopolyploidisation event – between *N. obtusifolia* and *N. sylvestris -* approximately 4-5 million years ago (Clarkson et al. 2005; Clarkson et al. 2017). The phylogenetic relationships among the species of this group have been thoroughly analysed based on nuclear and plastid DNA sequences, as well as using the abundance of repeats as systematic markers (Dodsworth et al. 2017), each time resulting in consistent results. Therefore, it constitutes a perfect study case to test the novel phylogenetic approach presented in the current work. Plant materials and Illumina sequencing details can be found in Dodsworth et al. (2017). The reads can be downloaded from NCBI Short Read Archive [SRA] using the following accession numbers: *N. repanda*—SRR453021; *N. nudicaulis*—SRR452996]; *N. sylvestris*—SRR343066; *N. obtusifolia*— SRR452993; *N. nesophila*—SRR4046065; *N. stocktonii*—SRR4046066. Illumina reads were trimmed to 95 nucleotides and quality filtered (quality score of 10 over 95% of bases and no Ns allowed) using the RepeatExplorer 2 pre-processing tools implemented in Galaxy. Then, 0.8 % of the genome proportion of each species was prefixed with a three-letter code unique to that species and combined to produce a dataset of 2,189,506 reads. The analysed genome proportion for *Nicotiana* species (as well as for the rest of the datasets) was determined according the available computing capacity of the RepeatExplorer Galaxy instance.

#### Fabeae

The legume tribe Fabeae represents a large and diverse group of angiosperms showing a long evolutionary history (Schaefer et al. 2012). Consequently, it represents an adequate case study to test the accuracy of our novel phylogenetic approach at higher taxonomic levels and analysing more distantly related taxa. In all species, genomic DNA was extracted from isolated leaf nuclei and sequenced on the Illumina platform (see Macas et al. 2015, for further details). All read data are available at the SRA with the following accession numbers: *Vicia hirsuta*—ERR413114; *V. ervilia*—ERR413112; *V. sylvatica*—ERR413113; *V. tetrasperma*—ERR413111; *Lathyrus sativus*—ERR413118 & ERR413119; *L. vernus*—ERR413116 & ERR413117; *L. latifolius*—ERR413120; *Pisum sativum*—ERR063464; *P. fulvum*—ERR413083. Illumina reads were pre-processed as in *Nicotiana* example, but in this case the sequences were trimmed to 90 nucleotides. Finally, 0.9 % of the genome proportion of each species was combined on a dataset of 4,943,339 reads.

#### Drosophila

In order to test our method on an animal study model, we focused on seven *Drosophila* species from the *melanogaster* subgroup. Illumina reads for the following species were downloaded from the SRA: *Drosophila bipectinata*—SRR345542; *D. suzukii*— SRR1002946; *D. biarmipes*—SRR345536; *D. ananassae*—SRR491410; *D. melanogaster*—SRR1005465; *D. sechellia*—SRR869587; *D. simulans*—SRR580349. Illumina sequences were pre-processed as explained above, with the reads trimmed to 76 nucleotides. In this case, given the small genome size of *Drosophila* species, 17 % of the genome proportion of each species was pooled in a dataset of 3,225,727 reads.

#### Asclepias

The Sonoran Desert clade of *Asclepias* represents a difficult phylogenetic problem to test the new phylogenetic approach we present here. The diversification process within this group is recent, encompassing some hybridization events (Straub et al. 2012). Consequently, the evolutionary reconstructions derived from plastid and ribosomal DNA showed important phylogenetic incongruence. In addition, genome size data for these *Asclepias* species is not available, so the number of input reads for the comparative clustering analysis could not be calibrated to include the same genome proportion of each sample. Potentially derived from this missing information, the phylogenetic signal obtained by Dodsworth et al (2014) using repeat abundances differed significantly from previous reconstructions based on plastid and ribosomal DNA. Illumina reads for species from the Sonoran Desert clade of *Asclepias* were downloaded from the SRA: *Asclepias macrotis* 149—SRX384308; *A. albicans* × *subulata* 282— SRX384307; *A. cutleri* 382—SRX384306; *A. subulata* 423—SRX384305; *A. macrotis* 150— SRX384304; *A. albicans* 422—SRX384303; *A. subulata* 411—SRX384302; *A. masonii* 154— SRX384301; *A. leptopus* 137—SRX384300; *A. cutleri* 421—SRX384299; *A. coulteri* 45— SRX384298; *A. subaphylla* 272—SRX384297; *A. subaphylla* 271— SRX384296; *A. albicans* 003— SRX384295. After pre-processing (following the same procedures than in the cases above), the reads of *Asclepias* species were trimmed to 70 nucleotides and the combined dataset was constituted by 1,349,066 reads.

### Clustering and phylogenetic analyses

Clustering of Illumina reads was performed using the RepeatExplorer 2 (RE2) pipeline, implemented in a Galaxy server environment (https://repeatexplorer-elixir.cerit-sc.cz) as described in Dodsworth et al. (2014). Separate comparative analyses (i.e. simultaneous clustering of reads from all species in the dataset) were run for each dataset on RE2, using default settings. Briefly, using a BLAST threshold of 90% similarity over 55% of the read length, RE2 identifies similarities between all sequence reads and then identifies clusters – representing homologous repeats – based on a principle of maximum modularity (Fig. 1b). Then, the most abundant repeats (i.e. constituted by more than 0.01% of input reads) are employed for phylogenetic analyses. Clusters identified as belonging to the plastid or mitochondrial genomes were removed prior to the phylogenetic analyses.

This novel method of phylogenetic reconstruction is based on the pairwise genetic distances between the repeatomes of the different species included in the datasets. RepeatExplorer 2 generates a first matrix showing the number of edges between species (i.e. the number of significant BLAST matches between the reads of the different species) for each of the most abundant identified clusters. The more similar the sequences of a repeat among two species, the higher the number of edges between the reads of those species (and vice versa). At the same time, RE2 calculates a second matrix with the observed/expected number of edges between species (Fig. 1c). The expected number of edges between two species is obtained by multiplying the total number of reads of these two species in the cluster and dividing by the total number of reads (of all the species) that constitute the same cluster. By calculating the observed/expected number of edges between species, the values of this second matrix are balanced by the number of reads that each species contributes to the cluster (i.e. the different repeat abundances between species). Therefore, the observed/expected number of edges between species is a measure of the pairwise similarity among the reads of the species (i.e. sequence similarity), which can be calculated for each of the most abundant clusters.

These similarity matrices are simply transformed to distance matrices by calculating the inverse of the values. Then, the function *NJ* from package *ape* (Paradis & Schliep, 2018) in R (R Core team, 2018) is employed to construct a neighbor-joining tree (based on distances) for each of the top 100 most-abundant clusters identified by RE2 (Fig. 1d). Incomplete matrices lacking pairwise similarity values (i.e. clusters/repeats that were not present in all species) were excluded. All these trees are exported to SplitsTree4 (Huson & Bryan, 2006) in Newick format to finally construct a consensus network (Holland & Moulton 2003) showing splits contained in more than 20% of the trees (Fig. 1e).

### Testing method performance

To test the performance of the method, several parameters were analyzed with the smallest clustering dataset (i.e. *Nicotiana* sect. *Repandae*), but using the modifications described below: i) To evaluate the number of clusters sufficient to obtain a resolved phylogenetic hypothesis, consensus networks were built from different numbers of clusters (i.e. top 10, top 50, top 100, and top 200), comparing the topology of the network inferred each time; ii) To evaluate the effect of weighting the number of reads for each species included in the comparative clustering in accordance to the genome size of the species, a network was inferred from a clustering analysis with equal amounts of reads (i.e. 350,000 reads) for all *Nicotiana* species. Then, the phylogenetic relationships among the species were compared between both networks; iii) Finally, the relative informativeness of different repeat types was analysed by creating subsets of the original matrix based on different repeat annotations. Annotations were assigned to the following categories based on BLAST hits to Repbase, custom-protein domain database in the RE2 pipeline and graph structure: DNA transposon, Ty1/*Copia* LTR retrotransposon, Ty3/*Gypsy* LTR retrotransposon, rDNA and satellite. Matrices were created based on each repeat type and networks were inferred as described above.

## Results

### Species relationships using repeat similarities

#### Nicotiana

After excluding those clusters lacking edges between reads of any pair of species, 85 neighbour-joining trees (out of the top 100 clusters) were constructed and summarized in the consensus network based on repeat similarities (Fig. 2a). The branches of the network showed no conflicting splits, indicating overall consistency in the topology of the individual trees employed to infer the phylogeny. The phylogenetic hypothesis we obtained was equivalent to previous ones based on nrITS, plastid markers, low-copy nuclear genes or repeat abundances (Chase et al. 2003; Clarkson et al. 2004; Kelly et al. 2013; Renny-Byfield et al. 2013; Dodsworth et al. 2017).

**Figure 2.**
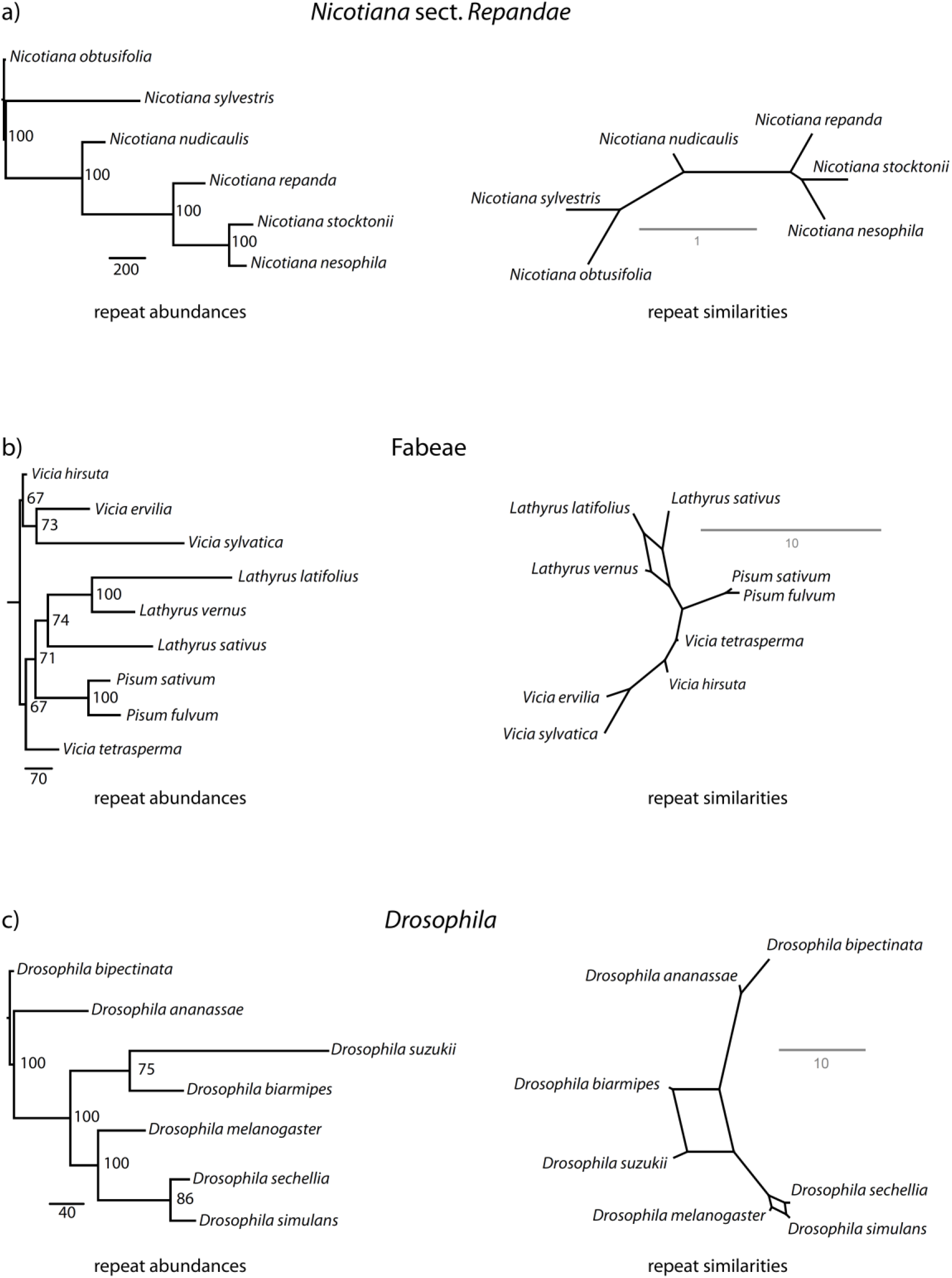
Phylogenetic relationships in *Nicotiana* sect. *Repandae*, Fabeae and *Drosophila* based on repeat abundances (left side; from Dodsworth et al. 2014, 2017) and repeat sequence similarities (right side; this study). Numbers on nodes in repeat abundances trees represent BPs ≥ 50 and scale bars show relative numbers of step changes. Scale bars from repeat similarities networks indicate mean branch length across all single cluster trees.

#### Fabeae

Out of the 100 most abundant clusters reconstructed by RE2, 40 matrices lacked pairwise similarity values between some of the species, so they were excluded from the phylogenetic analyses. In most cases, these clusters were only constituted by reads of a single species or genus, indicating that those repeats were probably exclusive of that particular taxonomic group. Using the remaining 60 complete matrices, the phylogenetic relationships inferred with our approach (Fig. 2b) were mainly the same than those obtained by Dodsworth et al. (2014) based on nrITS, plastid markers and repeat abundances. Our consensus network showed conflicting splits involving *Lathyrus* species, whereas repeat abundances and plastid approaches presented incongruent topologies and low statistical support, respectively, on these branches.

#### Drosophila

Also in this case, many clusters (63 out of the top 100 most abundant) inferred by RE2 were only constituted by reads from a part of *Drosophila* species. The matrices derived from these clusters lacked similarity values among some pairs of species, so they were discarded for the phylogenetic analyses. Consequently, only 37 clusters were employed to construct NJ trees and infer the consensus network (Fig. 2c). Despite being constructed from the similarity data of few repeats, this network still resembled the phylogenetic inferences based on nuclear DNA, mitochondrial DNA and repeat abundances (Dodsworth et al. 2014). The only differences affected the phylogenetic relationships within two groups (*D. suzuki–D. biarmipes* and *D. melanogaster–D. simulans–D. sechellia*), showing conflicting splits according to our results, while DNA sequences and repeat abundance data gave rise to highly resolved trees.

#### Asclepias

Out of the top 100 clusters reconstructed by RE2, 89 showed edges among the reads of each pair of species, so they were employed to infer neighbour-joining trees and subsequently summarized in the consensus network (Fig. 3). The phylogenetic relationships among *Asclepias* species inferred from this network showed comparable results with the evolutionary reconstructions of this group based on ribosomal DNA and plastomes (Dodsworth et al. 2014). As in those previous phylogenetic trees based on rDNA and cpDNA, our approach failed to reconstruct the species relationships within the hybridogenous group constituted by *A. albicans, A. masonii, A. subaphyla* and *A. subulata*. However, for the rest of *Asclepias* species included in the analysis, the consensus network based on repeat similarities inferred well their phylogenetic relationships. Mirroring as well the evolutionary inferences based on rDNA and plastid DNA, our approach indicates that *A. coulteri* and *A. macrotis* constitute a monophyletic group, while the systematic placement of *A. cutleri* and *A. leptolopus* is not fully resolved. In contrast, in this study case, the reconstruction based on repeat abundances (Dodsworth et al. 2014) showed a completely different hypothesis. As proposed by the same authors, their results could be potentially compromised by a lack of genome size data with which to calibrate the number of input reads.

**Figure 3.**
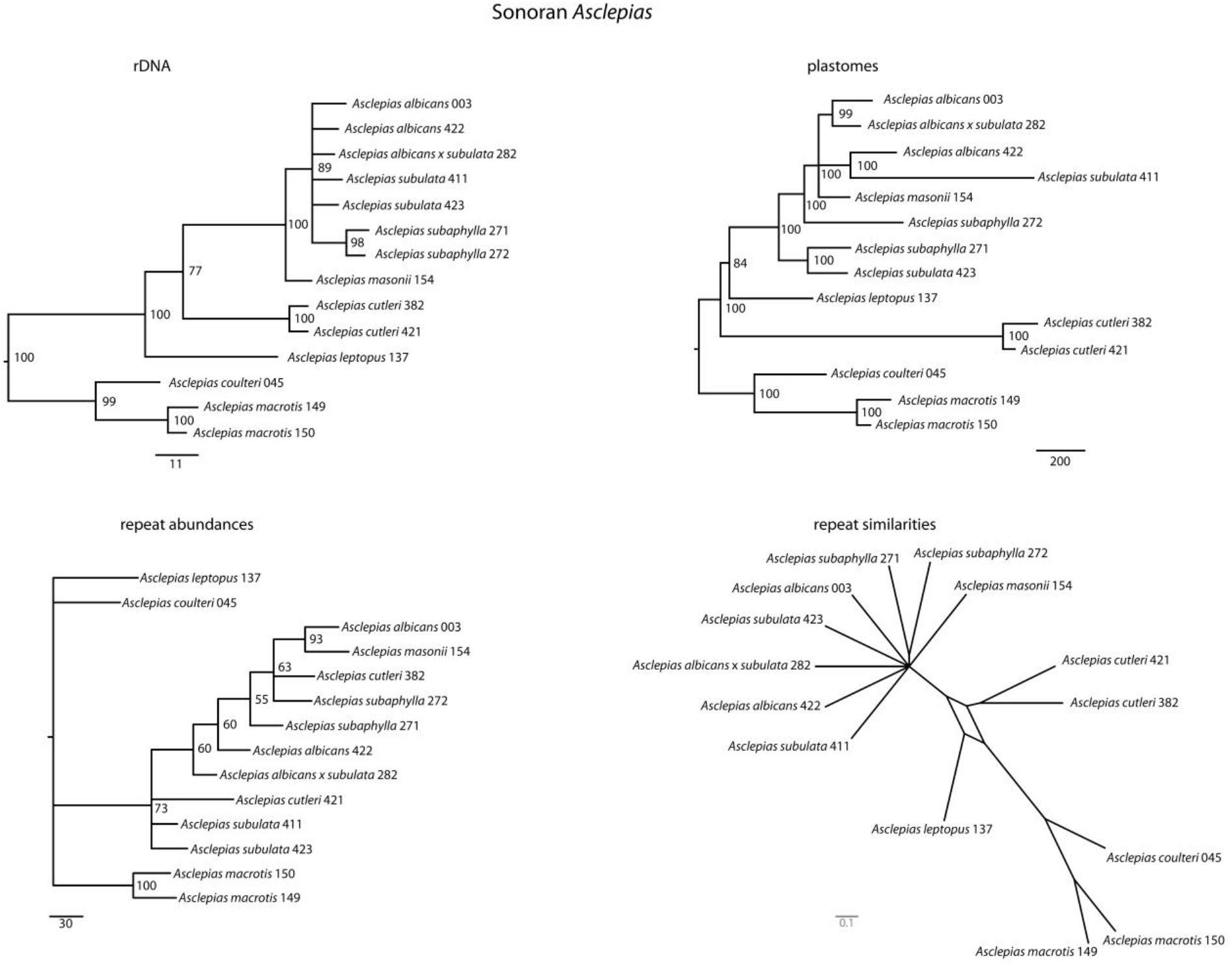
Phylogenetic relationships in Sonoran *Asclepias* taxa based on rDNA, plastomes and repeat abundances (from Dodsworth et al. 2014) as well as repeat sequence similarities (this study). Numbers on nodes in rDNA, plastome and repeat abundances trees represent BPs ≥ 50 and scale bars show relative numbers of step changes. Scale bars from repeat similarities network indicate mean branch length across all single cluster trees.

### Evaluating method performance

According to our analyses with the *Nicotiana* dataset, the subset of clusters producing a network that better resolves the phylogenetic relationships among species is top 100 (Fig. 4c). With lower numbers of clusters (e.g., top 10 or top 50), the network retains considerable phylogenetic information, but some inconsistency and lack of resolution appears within the group constituted by *N. repanda, N. stocktonii* and *N. nesophila* (Fig. 4a, c). Considering higher number of clusters (i.e. the 200 most abundant), the phylogenetic resolution of the network decreases (Fig. 4d), indicating that less abundant clusters show inconsistent systematic information that swamps the true phylogenetic signal. In addition, the number of matrices lacking pairwise similarity values (therefore not considered to construct NJ trees) become much more frequent among the clusters 100 to 200 (50 clusters) than in the first 100 clusters (15 clusters). This is likely due to lower-abundance repeats tending to show more species-specific patterns.

**Figure 4.**
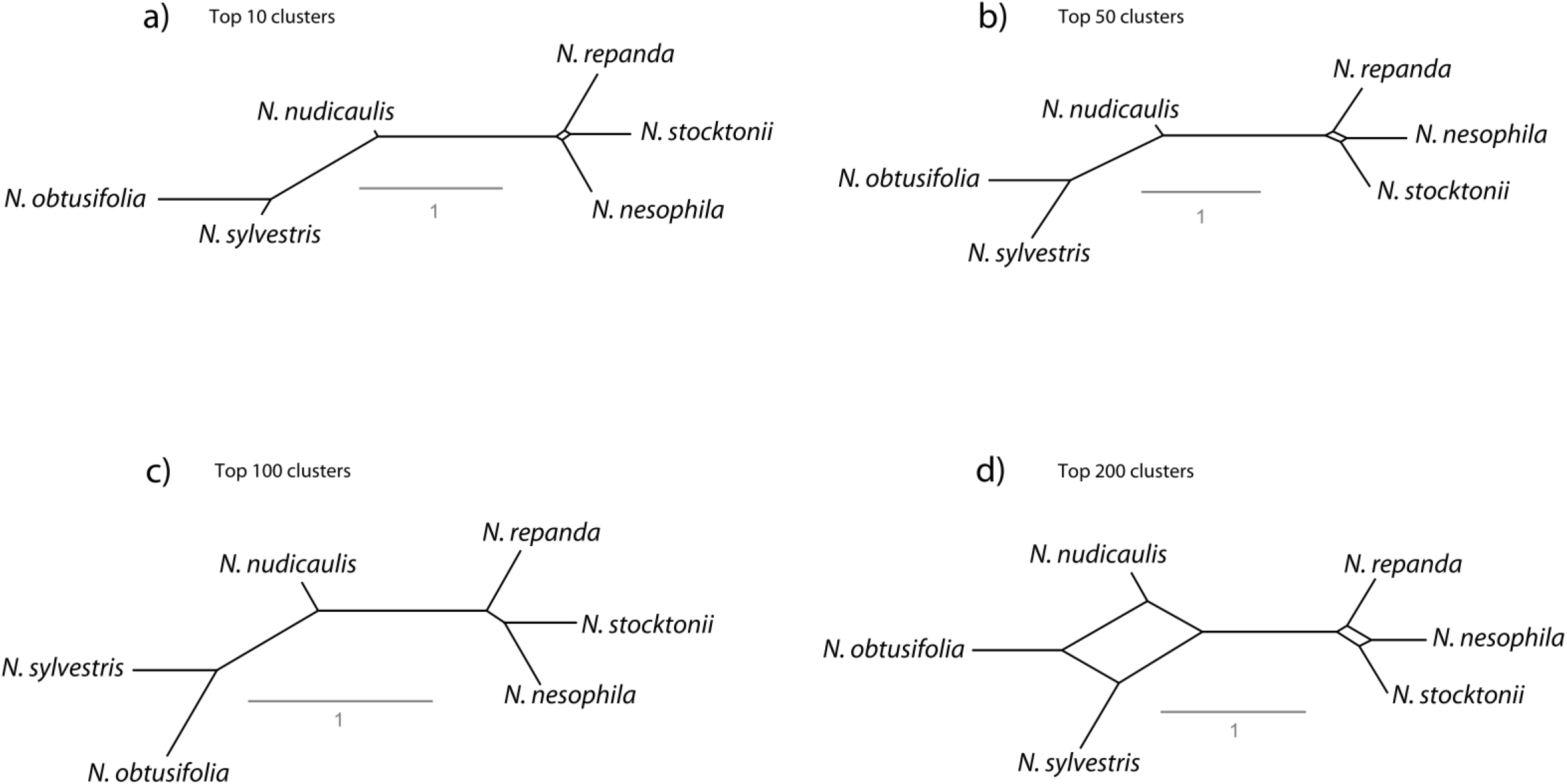
Performance test of the cumulative number of clusters (a, top 10 clusters; b, top 50 clusters; c, top 100 clusters; d, top 200 clusters) employed to infer the phylogenetic relationships in *Nicotiana* sect. *Repandae*. Scale bars from repeat similarities network indicate mean branch length across all single cluster trees.

The result of our test evaluating the effect of including the same number of reads per species (instead of weighting the number of reads according to the genome size of the samples) can be observed in Fig. 5. Without accounting for genome size differences (Fig. 5b), the output network shows some conflicting splits within the *N. repanda, N. stocktonii* and *N. nesophila* group, but no evidence of spurious relationships was found. The phylogenetic relationships among the rest of the species is the same as in the result obtained after resampling the number of reads in relation to genome sizes (i.e. same genome proportions).

**Figure 5.**
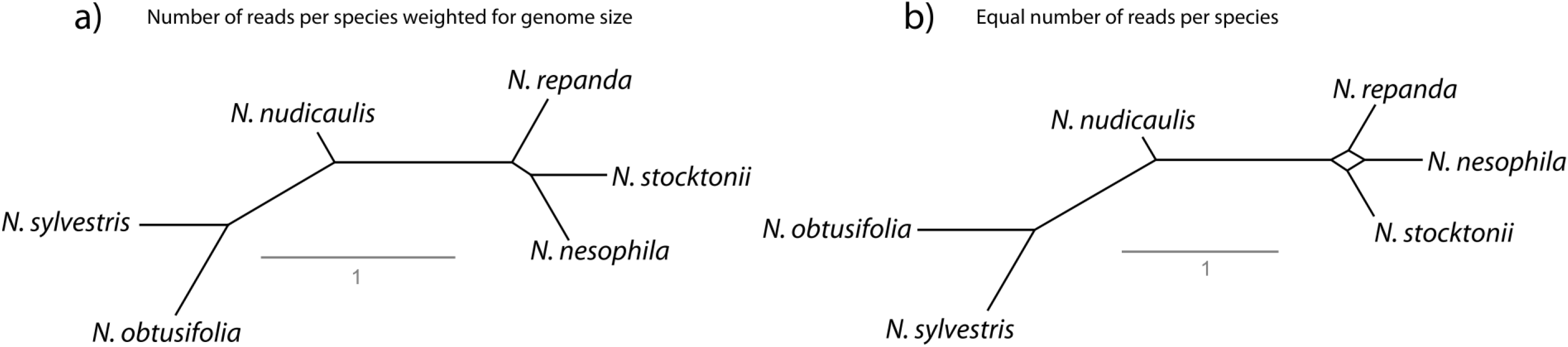
Performance test of the relative number of reads per species (a, number of reads weighted for genome size; b, equal number of reads per species) employed to infer the phylogenetic relationships in *Nicotiana* sect. *Repandae*. Scale bars from repeat similarities network indicate mean branch length across all single cluster trees.

The performance of the method with respect to the variety of repeat types characterized by RE2 is shown in Figure 6. The relative informativeness of the different classes of repeats appears to be related to their abundance in the genome. In this way, the most abundant repetitive elements in *Nicotiana* genomes (i.e. Ty3/*Gypsy* LTR repeats, contributing 34 clusters to the network) produced a phylogenetic reconstruction perfectly mirroring the network obtained from all the repeats. In contrast, the network inferred from Ty1/*Copia* LTR elements (accounting for 16 clusters in the analysis) showed some splits within the *N. sylvestris, N. nudicaulis* and *N. obtusifolia* group. Other less abundant repeat types (e.g. DNA transposons, 11 clusters; ribosomal DNA, 4 clusters) also produced evolutionary informative networks. However, in the case of satellites (showing only two clusters constituted by reads from all the species), the low abundance may have compromised phylogenetic resolution.

**Figure 6.**
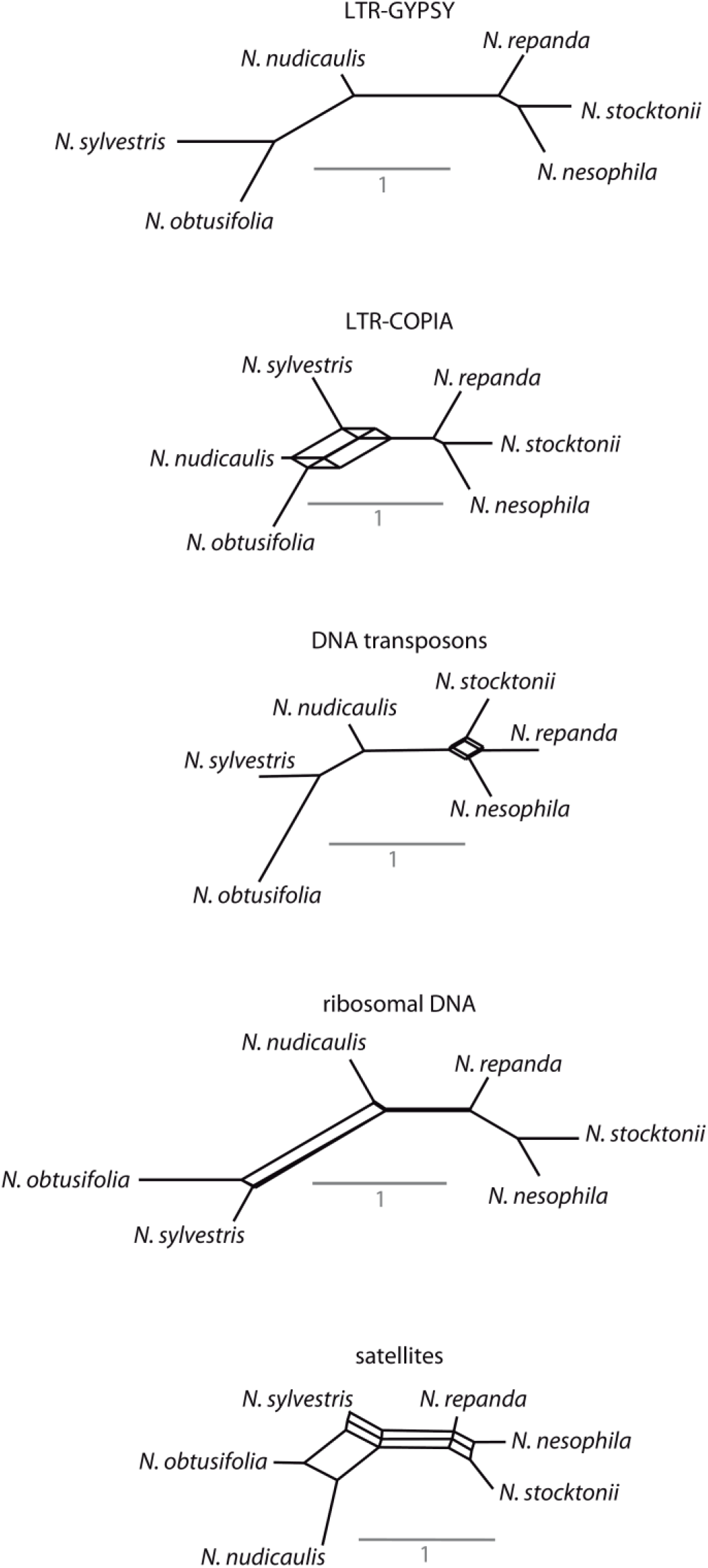
Performance test of repeat types employed to infer the phylogenetic relationships in *Nicotiana* sect. *Repandae*. Scale bars from repeat similarities network indicate mean branch length across all single cluster trees.

## Discussion

The results presented in this study show that pairwise read similarities between homologous repetitive elements can be employed to resolve species relationships. The consensus networks inferred from the analyses of *Nicotiana*, Fabeae, *Asclepias* and *Drosophila* repeatomes showed mainly defined topologies, indicating overall consistency in the structure of the individual trees employed to construct the networks. The phylogenetic relationships we obtained mirror previous evolutionary reconstructions of these groups found with more standard sequence markers and analyses. Our results also suggest that the method is useful to infer phylogenetic relationships at considerably different taxonomic levels, as long as the members of each group share enough homologous repeat clusters. Even employing similarity data from a low number of repeats (e.g. less than 50 clusters), the networks we obtained reveal significant phylogenetic signal. Furthermore, the informativeness of the repeats seems to be related to their relative abundance but not to the repeat categories they represent, thus the utility of this approach should be robust to the differences in repeatome composition of a wide variety of organisms.

Except for the *Asclepias* dataset, the phylogenetic analyses based on repeat sequence similarities produced virtually the same hypotheses as those obtained by Dodsworth et al. (2014; 2017) based on repeat abundances. These results indicate that, at least for these study cases, repeat sequence divergence and repeat abundance may evolve in concert, most probably following the process of random genetic drift (Jurka et al. 2011; 2012). In the case of *Asclepias*, the trees inferred from the repeat abundances approach (Dodsworth et al. 2014) showed conflicting phylogenetic relationships compared to those obtained from plastid and rDNA sequence data. Indeed, the same authors advised that their systematic hypothesis on this group could be biased by the lack of genome size data. Conversely, the results we obtained with repeat sequence similarities depicted an evolutionary history fully concordant with previous more standard phylogenetic studies of *Asclepias* (Straub et al. 2012). Therefore, our study reinforces the idea that genome size calibration is essential to produce reliable phylogenetic results using repeat abundances, whereas the repeat sequence similarities approach appears to be more robust in the absence of genome size data.

These phylogenetic approaches using repeat abundances and repeat sequence similarities should be seen as complementary to infer species relationships based on repeatomic data. Moreover, the joined application of both strategies could be suitable to resolve particular evolutionary questions linked to the divergence of the repeatome. We foresee that studying both the evolution of repeat abundance and sequence divergence on the same datasets may help better understand the mode of evolution of genomes. The combined use of both approaches could be helpful to reveal scenarios where repeatome evolution does not obviously follow simple random genetic drift processes. For instance, chromosome reorganizations (e.g. segmental insertions, deletions and/or translocations) are supposed to involve sudden changes in genome and repeat abundance, contrasting to more gradual processes such as transposable element accumulation or tandem repeat amplification over time (Schubert & Vu 2016). In *Anacyclus*, genomic evolution, likely driven by chromosomal rearrangements, induced changes in repeat abundances that do not correspond with repeat sequence divergence (Vitales et al., in prep.). Indeed, repetitive DNAs have been proposed to be key elements for understanding the mechanism and dynamics of chromosomal rearrangements among eukaryotic genomes (Eichler & Sankoff, 2003). Therefore, the comparative evolutionary signal of repeat abundances and repeat sequence similarities could be indicative of the mechanisms governing genome evolution. The increasing availability of genome skimming data could extend these repeatome analyses to many species. Additionally, cytogenetic information on processes involving chromosome restructuring could help us understand sudden changes in repeatome evolution as well as the potential impact on its intrinsic phylogenetic signal.

Finally, we propose that further analytical methods could be developed to solve some of the problems envisaged in this work. For instance, most distance-based phylogenies come with bootstrap support values, which are computed by resampling with replacement columns of homologous residues from the nucleotide alignment (Felsenstein, 1985). Unfortunately, this method cannot be applied to our repeat distances matrices, as they are based on pairwise read similarities. The problem is comparable to that faced by phylogenetic approaches based on pairwise distances between genomes obtained from alignment-free methods (e.g. Haubold, 2013; Ulyantsev et al. 2016; Zielezinski et al. 2017). However, the recent development of methods to estimate branch support from other kinds of pairwise distance data (e.g. Klötzl & Haubold 2016) look promising and could be applied in the future to the phylogenetic approach presented here. Regarding the clustering and similarity matrix construction, our method also shows considerable opportunities for fine-tuning. At present, this approach has only been tested by using the default pairwise BLAST comparison threshold of 90% similarity over 55% of the read length implemented in RE2 pipeline. This may limit the current approach to lower phylogenetic levels, however the modification of these parameters could have the potential to compare repeatome similarity at different evolutionary and taxonomic levels. This would permit the detection of enough homologous repeat clusters to perform phylogenetic reconstruction in very different study cases. In summary, the full possibilities of repeat abundance and sequence similarity are yet to be realised in comparative studies investigating genome evolution.

## Acknowledgments

We would like to thank Oriane Hidalgo, Teresa Garnatje, Jiri Macas and Petr Novak for insightful comments on previous versions of this manuscript. This work has been supported by DGICYT (Spanish Government; projects CGL2013-49097-C2, CGL2016-75694-P and CGL2017-84297-R) and the Generalitat de Catalunya (“Ajuts a grups de recerca consolidats” 2017SGR01116). S.G. benefitted from a Ramón y Cajal postdoctoral contract (RYC-2014-16608) from the government of Spain.

